# Latent periodic process inference from single-cell RNA-seq data

**DOI:** 10.1101/625566

**Authors:** Shaoheng Liang, Fang Wang, Jincheng Han, Ken Chen

## Abstract

Convoluted biological processes underlie the development of multicellular organisms and diseases. Advances in scRNA-seq make it possible to study these processes from cells at various developmental stages. Achieving accurate characterization is challenging, however, particularly for periodic processes, such as cell cycles. To address this, we developed Cyclum, a novel AutoEncoder approach that characterizes circular trajectories in the high-dimensional gene expression space. Cyclum substantially improves the accuracy and robustness of cell-cycle characterization beyond existing approaches. Applying Cyclum to removing cell-cycle effects leads to substantially improved delineations of cell subpopulations, which is useful for establishing various cell atlases and studying tumor heterogeneity. Cyclum is available at https://github.com/KChen-lab/cyclum.

## Background

Convoluted biological processes, which involve cell proliferation, differentiation, state transition, and cell-to-cell communication [1,2]. The course of development can be influenced by genetic (e.g., mutations), epigenetic, and environmental factors. Alterations to the genome, transcriptome, and proteome of individual cells also can result in pathogeneses [3]. Early efforts have been made to reconstruct the temporal ordering of biological samples using bulk data [4,5], although challenges associated with cellular heterogeneity make it difficult to infer accurate time series. Advances in single-cell RNA sequencing (scRNA-seq) enabled large-scale acquisition of single-cell transcriptomic profiles and provided an unprecedented opportunity to uncover latent biological processes that orchestrate dynamic expression of genes in single cells throughout the course of the development [6]. However, it is very challenging to deconvolute these processes from scRNA-seq data accurately. A sufficiently large number of cells across time, lineage, and space need to be sampled in order to capture detailed sub-populational features and reduce technological noise. Tremendous efforts have been made to develop trajectory inference methods from scRNA-seq data. Over 59 methods have been developed since 2014 [7], including the widely known Monocle and Wanderlust. These methods represent biological processes in linear, bifurcating, or other graph topologies.

In many developmental processes, such as embryogenesis, organogenesis, and tumorigenesis [8], cell cycle plays a fundamental role. Distinct from processes that evoke linear changes in gene expression, cell cycle causes periodicity. A cycle starts from the G1 phase, goes through S and G2/M, and then returns to G1 within 24-hours for human cells [2]. This process is orchestrated elegantly by variable sets of genes (e.g., cyclins and cyclin-dependent kinases) that are turned on and off at relatively precise timings. As a result of such periodicity, the cycling cells at different transcriptomic states form a circular, non-linear trajectory in high-dimensional gene expression spaces. The positions of a cell alongside the circular trajectory indicate its timing (pseudo-time) in the cell cycle. Although well regulated, the process can be stochastic. For instance, cells can experience different fates (e.g., going into apoptosis or senescence), and the rate of development may fluctuate due to endogenous or exogenous factors [9].

Existing trajectory/pseudo-time inference methods are not optimal for representing such nonlinear periodicity. Those based on linear representations, such as principal component analysis (PCA), cannot accurately represent circular timings or infer effect sizes. The cell cycle regression approaches implemented by scLVM [10], Seurat [11], and ccRemover [12] are based on linear representations generated from user-defined gene sets, which may be biased or incomprehensive, particularly in cancer cells with aberrant cell cycle. These regression methods do not explicitly calculate pseudo-time, which limits their utility in data analysis. Cyclone [13] uses PCA and relative expression of gene pairs to predict cell-cycle phases, which appears to perform better than traditional machine learning methods, such as random forest, logistic regression, and support vector machine (SVM). A recent method reCAT [14] reconstructs cell-cycle pseudo-time using a Gaussian mixture model (GMM) to cluster single cells into groups and a quasi-optimal traveling salesman path (TSP) solver to order the groups. The resulting pseudo-time is expressed in consecutive integers indicating the order of cells, instead of continuous real-number timings of the cells. Neither Cyclone nor reCAT can be applied to remove cell-cycle effects from the expression data.

To address these limitations, we developed an *ab initio* inference method, namely Cyclum, which employs a novel Auto-Encoder approach to capture the circular trajectory in the high dimensional gene expression space, formed by single cells sampled from various stages of a periodic process. Conceptually, our approach identifies an optimal (least square) embedding of cells in a circular space described by periodic kernel functions. It effectively unfolds the circular manifold onto a linear space to obtain precise pseudo-time (Figure 1 and Additional file 1, Figure S1). During the preparation of this manuscript, Saelens et al. [15] introduces a method to infer pseudo-time only from the first two principal components. Cyclum constructs the pseudotime in a more flexible way.

**Figure 1:**
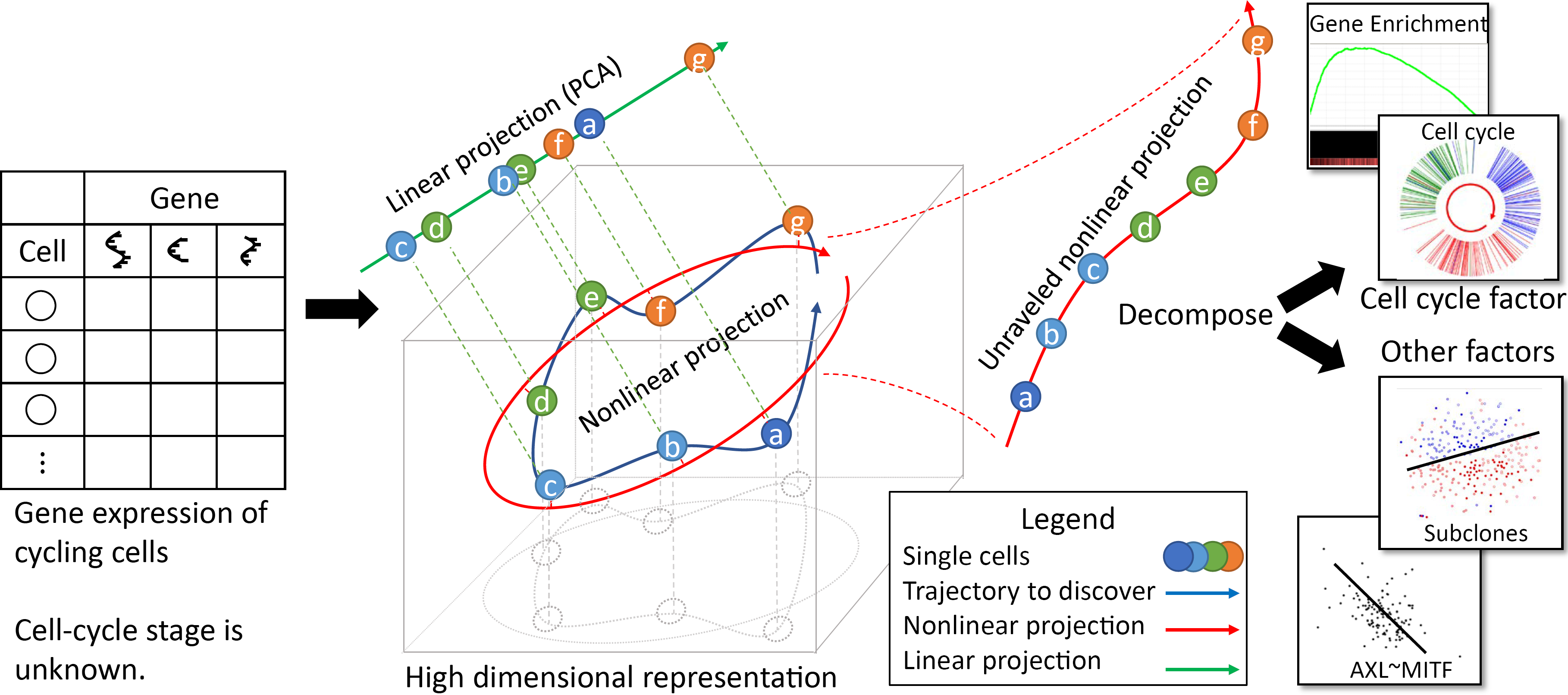
Overview of the Cyclum program. Single-cell RNA-seq data in the format of a cell-gene expression matrix are given to Cyclum, which identifies a circular trajectory consisting of cells at different times (indicated by English letters) and stages (labeled by colors) in the high-dimensional gene expression space. Cyclum unravels the circular trajectory (red arrows) along with the projected cells to infer their pseudo-time. In contrast, a linear projection (green arrow) would result in incorrect ordering and timing. The inferred genes and pseudo-times can be further analyzed to discover new functions, cell-types, and cell-phenotype associations.

## Results

### Overview of Cyclum

In a nutshell, the Cyclum program (Figure 1) analyzes a cell-gene expression matrix using an Auto-Encoder technique (see Methods), which projects the cells onto a non-linear periodic trajectory, where the pseudo-time of the cells in a periodic process can be more accurately determined than with linear approaches, such as PCA. Cyclum can be used to identify genes associated with the periodic process, based on the degree of match between the kinetics of gene expressions and the inferred periodicity. Additionally, this program can treat the inferred periodic process as a confounder and deconvolute its effects from scRNA-seq data. Using Cyclum in this way can result in enhanced delineations of cell subpopulations segregated by lineages or phenotypes.

### Accuracy of Cyclum for cell-cycle characterization

We compared Cyclum’s performance for characterizing cell-cycles with Cyclone, reCAT, and PCA. Four datasets were used (Table 1). The first dataset was obtained from a set of mouse embryonic stem cells (mESC) [10] using SMARTer kit and Illumina HiSeq 2000 sequencing technology. The other three datasets were obtained using nanoString nCounter technologies from the bone metastasis of a prostate adenocarcinoma (PC3), the pleural effusion metastasis of a breast adenocarcinoma (MB), and the lymphoblast node of a lymphoma (H9) [16]. Each of these datasets has 200 to 400 cells. Flow sorting with Hoechst staining was performed on the same set of cells, and the cells were then classified into three stages G0/G1, S, and G2/M, based on their DNA mass.

**Table 1.** Cell-cycle scRNA-seq datasets.

| Dataset | Assaying | Labeling | Cell count |  |  |  | Gene count |
| --- | --- | --- | --- | --- | --- | --- | --- |
|  |  |  | Total | G0/G1 | S | G2/M |  |
| mESC | scRNA | Hoechst | 288 | 96 | 96 | 96 | 38,293 |
| PC3 (CRL-1435) | qPCR | Hoechst | 361 | 85 | 141 | 135 | 253 |
| MDA-MB-231 (HTB-26) | qPCR | Hoechst | 342 | 123 | 103 | 116 | 253 |
| H9 (HTB-176) | qPCR | Hoechst | 227 | 66 | 68 | 93 | 253 |

We ran each algorithm on each dataset and classified cells into various cell-cycle stages. Discretization of the continuous Cyclum and reCAT results was accomplished using a three-component Gaussian mixture model. We then calculated the fraction of cells that were correctly classified by comparing the predicted cell-cycle labels with those obtained from the flow-sorting. As shown (Figure 2a), Cyclum outperformed for all four datasets vs. the other four methods, including Cyclone and reCAT, which used known cell-cycle genes to optimize their performances. We also performed Gene Set Enrichment Analysis (GSEA) [17,18] of the genes discovered *ab initio* using Cyclum, PC1, and PC2 on the mESC dataset. Cyclum yielded a normalized enrichment score (NES) for cell-cycle genes of 1.57, which compared favorably with the PC1 and PC2 scoring of 1.07 and 1.06, respectively, showing that Cyclum can better infer cell-cycle genes than any of the principal component analyses.

**Figure 2:**
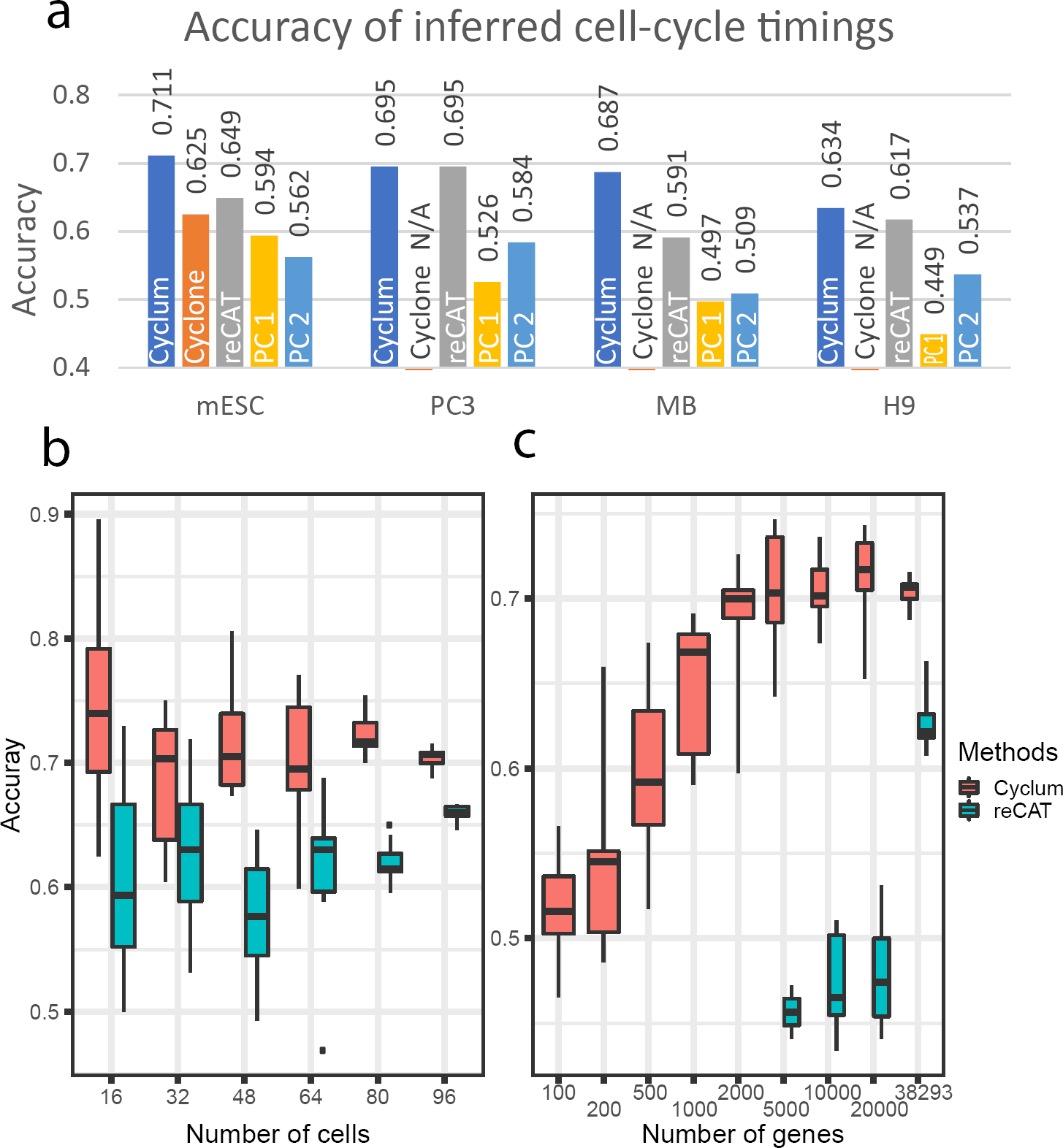
Accuracy of cell-cycle inference. **a)** Cell cycle classification accuracy of Cyclum compared with Cyclone, reCAT, and principal component 1 (PC1) and 2 (PC2). **b)** Cell-cycle classification accuracy (Y-axis) of Cyclum (red) and reCAT (green) over the number of cells (X-axis). Plotted are the range (box) and the mean (horizontal bar) of the results obtained from 10 randomly down-sampled datasets. **c)** Cell-cycle classification accuracy (Y-axis) over the numbers of genes (X-axis, log-scale). Plotted are the range (box) and the mean (horizontal bar) of the results obtained from 10 randomly down-sampled datasets. The numbers on the boxes are the times of successful reCAT runs.

We further assessed the robustness of Cyclum as related to data sparsity. We randomly down-sampled the mESC data for fewer cells or genes. Stratified subsampling was used to keep an equal number of cells in each stage. We observed that the mean classification accuracy of Cyclum (ranging between 0.7 and 0.75) remained largely invariant with regard to the number of cells. In contrast, the mean accuracy of reCAT became substantially worse with fewer cells (Fig. 2b). The variance increased with fewer cells for both programs. In a parallel experiment, we uniformly randomly down-sampled genes. The mean accuracy of Cyclum was unaffected when there were over 10,000 genes (Figure 2c). The performance of reCAT was substantially worse, however, with fewer genes, and a failure to return results when there were less than 5,000 genes.

### Separability of subclones after corrected for cell cycle

We assessed the utility of Cyclum in reducing the confounding effects introduced by cell-cycling. A tissue sample often consists of multiple types of cells (e.g., tumor subclones) with distinct transcriptomic profiles [1,19]. When the cells are under active cycling, it can become difficult to delineate the cell types.

To assess the utility of Cyclum in this setting, we generated a virtual tumor sample consisting of two proliferating subclones of similar, but different transcriptomic profiles. We used the mESC data as one clone, and we created a second clone by doubling the expression levels of a randomly selected set of genes containing variable numbers of known cell-cycle and non-cell-cycle genes (see Methods). We then merged cells from these two clones together as a virtual tumor sample. This strategy allowed us to use real scRNA-seq data, although the perturbations applied are artificial. More importantly, it allowed us to track the clonal origins of each cell in the mixed population. We then ran Cyclum, ccRemover, Seurat, and PCA on the virtual tumor samples created under a wide range of parameters and assessed the accuracy of the algorithms in delineating cells from the two subclones. Cyclone and reCAT cannot remove cell-cycle effects, thus they were not included in the assessment.

We found that cells from the two subclones in a virtual tumor sample are intermingled in the tSNE plot that was generated from the scRNA-seq data (Fig. 3a). After removing cell-cycling effects using Cyclum, cells in the two subclones became separable (Fig. 3b). We then performed systematic assessment under a range of parameters, including the number of cells, number of perturbed genes, and the fraction of cell-cycling genes. We used two-component Gaussian mixture models to quantify how well the two subclones were separated (classification accuracy) in the t-SNE plot. Under almost all conditions, Cyclum achieved significantly higher accuracy than the other methods, particularly when a large number (>400) of cell-cycling genes were perturbed (Figure 3c and Additional file 1: Figure S2). In contrast, approaches such as Seurat and ccRemover, which rely on known cell-cycling genes, performed worse, especially when more cell-cycling genes were perturbed. These results demonstrated the benefit and robustness of Cyclum in deconvoluting cell-cycling effects from the scRNA-seq data.

**Figure 3:**
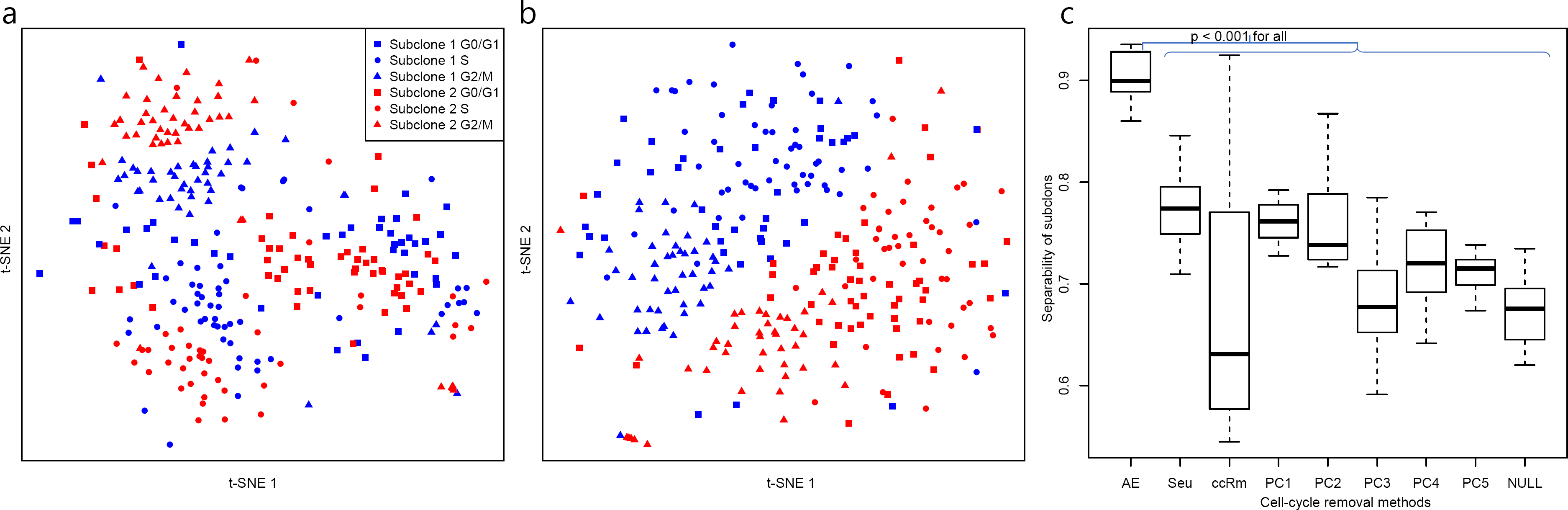
Subclone detection from virtual tumor data. **a)** t-SNE plot of the virtual tumor data consisting of two subclones (blue and red dots) of 288 cells each at various cell-cycling stages. **b)** t-SNE plot of the data corrected for cell-cycling effects using Cyclum. **c)** The range (box) and the mean (horizontal bar) separablility (Y-axis) over 10 randomly generated virtual tumor datasets using data corrected by Cyclum, Seurat (Seu), ccRemover (ccRm), and Principal Component (PC) 1~5, and the uncorrected data (NULL). The expression levels of 1,600 genes, including 600 known cell cycling genes that were doubled in creating the virtual tumor data. P-values were calculated using two-side Student’s t-test.

### Application of Cyclum on the melanoma data

We further examine the utility of Cyclum in analyzing scRNA-seq data obtained from real cancer samples. We examined the dataset [20] consisting of the RNA expression of 23,686 genes in 4,645 single cells from 19 melanoma patients, profiled using the 10X Chromium technology.

We analyzed the data from the five patients (i.e., 78, 79, 80, 81, and 88) that had over 100 cancer cells. First, we assessed how accurately Cyclum could discover cell-cycle genes (see Methods). We compared the pseudo-time inferred by Cyclum, reCAT, PC1, and PC2 against the GO:0007049 GO_CELL_CYCLE gene set using the GSEA. A higher GSEA score indicates that the pseudo-times inferred are more accurately tracing cell cycle. Cyclum performed the best in this analysis (Figure 4a and Additional file 1, Table S2), even on samples that reportedly had few cycling cells (e.g., Mel79). Among the novel cell-cycling genes nominated by Cyclum (Additional file 1, Table S2, Figure S3, and S4), *KCNQ1OT1* and *FBLIM1* (Additional file 1, Figure S3, and Table S3) have recently been shown in the literature to be related to proliferation and tumorigenesis [21–26].

**Figure 4:**
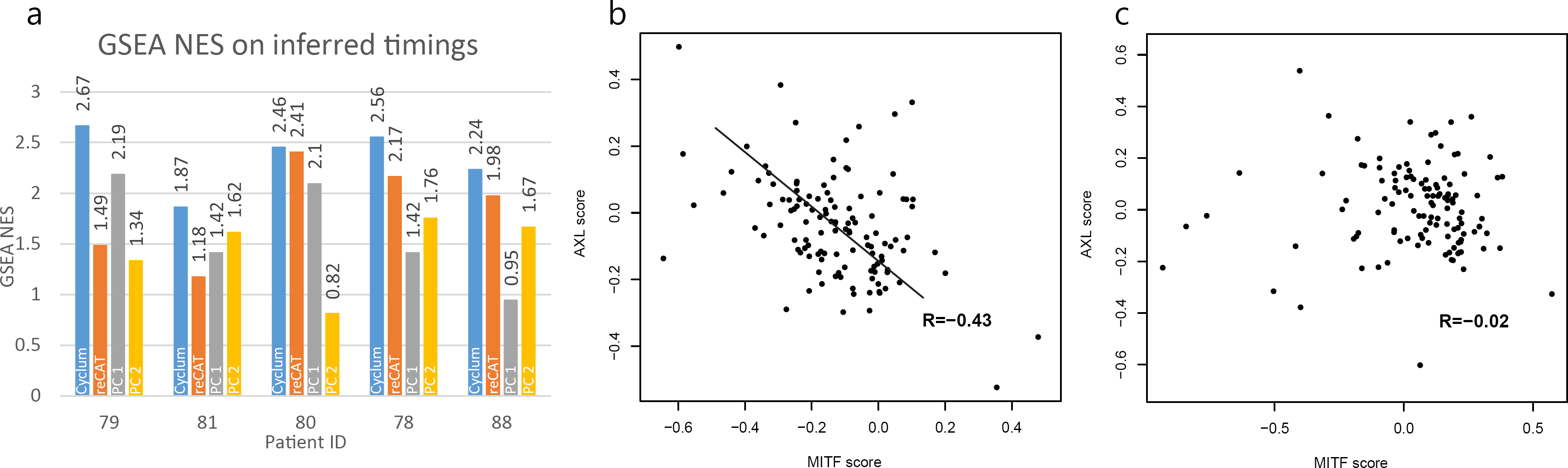
Cyclum results on the melanoma data. **a)** GSEA NES scores were obtained based on pseudo-times inferred by Cyclum, reCAT, PC1, and PC2. **b)** The correlation between the MITF and the AXL scores for sample Mel78, based on Cyclum corrected expression data; **c)** uncorrected expression data. The AXL and MITF scores were calculated based on the average expression levels of the reported AXL and the MITF genes [20]. The line in (b) was drawn manually for visual reference. The R values are the Pearson correlation coefficients.

We estimated the proportions of cycling cells in these samples using Cyclum. Although Cyclum does not directly model quiescent cells, samples with fewer cycling cells (e.g., MEL79) appeared to have large gaps in the inferred pseudo-times (Additional file 1, Figure S5). These gaps corresponded well to the missing S, G2 and M stages in these samples. In contrast, samples with a large fraction of cycling cells (e.g., mESC) had largely continuous pseudo-times. By partitioning the pseudo-time densities, we estimated the fraction of cycling cells in each sample (Additional file 1, Table S1). The resulting fractions appeared consistent, but were generally higher than those estimated based on the expressions of marker genes [20]. That could be expected, as Cyclum summarized contributions from a larger set of genes showing periodicity.

Two dormant drug resistance programs (MITF-high and AXL-high) were present mutually exclusively in these melanoma patient samples, based on the immunofluorescence staining data [20]. To calculate the AXL/MITF program scores, we followed the method and gene sets suggested in [20]. The scores were defined as the average expression of the sets of genes. However, cell-cycling could confound the expression profiles of these cells [12], making it difficult to delineate the resistance subgroups. Indeed, before correcting for cell-cycling effects, almost no correlation (Figure 4c, *R* = −0.02, *p* = 0.81) was observed between the expressions of the cells from the two mutually exclusive programs in the actively proliferating sample Mel78. After applying Cyclum correction, a clearly negative correlation (Figure 4b, *R* = −0.43, *p* = 9 × 10^−7^) emerged, which is consistent with the expected mutual exclusivity between the two programs in single melanoma cells. The result was also better than that obtained using ccRemover and Seurat (Additional file 1, Figure S6).

## Discussions

In this work, we developed a novel trajectory inference method that can effectively characterize latent periodic developmental processes, such as cell cycles from scRNA-seq data. Compared with currently available methods, most of which are based on linear representations of the data, our approach can more effectively capture the non-linearity and achieve more accurate characterization of a periodic process.

We examined Cyclum using multiple real and synthetic datasets. Using cancer cell lines and mouse embryonic stem cell data, we demonstrated that Cyclum accurately infers cell-cycle timing from the gene expression profiles of single cells, which is validated by flow-sorting results obtained independently on the same set of cells. Using virtual tumor data, we showed that Cyclum can be applied to remove confounding cell-cycle effects and achieving an improved classification of distinct cell subpopulations. Although virtual tumor information cannot fully replace real data, the inputs were created under a wide range of parameters that facilitated systematic assessment of Cyclum and other comparable programs. Using the real datasets obtained from melanoma patients, we showed that Cyclum can accurately infer the cell-cycle expression components, nominate novel cell-cycle genes (e.g., *KCNQ1OT1* and *FBLIM1*), and elucidate latent associations between cell subpopulations and drug resistance. These experiments indicated that Cyclum can be applied as a generic tool for characterizing periodic processes and discovering biologically meaningful cell subpopulations from scRNA-seq data.

We anticipate that Cyclum will be able to impact several important areas of investigation. First, it may be applied to discovering new genes involved in a periodic process, particularly genes that have a transitional or relatively low expression, and whose relevance is only evident when being observed across time. Second, it can be applied to remove cell-cycle effects and enhance the characterization of cell types and developmental trajectories. These utilities will be in great demand by the Human Cell Atlas [27], the Human Tumor Atlas Network [28], and many other projects.

It is worth noting that Cyclum is a model-based approach that fits the data to predefined circular manifolds. This design makes Cyclum more robust to handle random noise and small sample sizes. This is a tremendous advantage over other model-free approaches, such as reCAT, for the purpose of characterizing cell cycles. Evidently, Cyclum’s demonstrated robustness to a reduced number of cells and genes makes it desirable to analyze current scRNA-seq datasets, which often suffer from cell-specific dropout and amplification bias [29]. Cyclum also appeared to work better on data that was heavily confounded by cell cycles. This is an important feature for studying cancer data, as many cancer cells have heightened cell-cycling activities [30,31].

On the other hand, when the latent process does not fit the circular manifold well, the method may or may not bring any benefit. Nonetheless, our study clearly demonstrated the advantage of fitting scRNA-seq data to circular manifolds in a variety of settings. We plan to further explore how to use Cyclum in conjunction with other methods to deconvolute data generated by more complex, intertwined processes. For example, we plan to explore Gaussian Process Latent Variable Models (GPLVM) to track a generic periodic manifold that is not restricted to a sinusoid in the high-dimensional expression space. GPLVM has been applied previously to model linear trajectories, but also potentially can be expanded to model periodic trajectories. We plan to investigate the potential of applying Cyclum to characterize other periodic processes, such as circadian rhythms [32], as well.

This work also demonstrates that unsupervised machine-learning techniques, such as AutoEncoders, can be successfully applied to model latent periodic processes, with innovations on the network architecture and activation functions. The Cyclum package is efficiently implemented in Python using TensorFlow [33], and it has been comprehensively tested. For example, Cyclum can analyze an scRNA-seq dataset consisting of 480 cells (23,686 genes) in a laptop computer with a GTX 960M GPU and 2 GB graphic memory in 10 minutes. Running without GPU on an Intel i7 5700HQ, a 3.5 GHz CPU returned the results in one hour. We anticipate that Cyclum can be easily scaled up to bigger datasets in a high-performance computer cluster.

## Conclusions

We developed Cyclum, a machine learning approach that can effectively and efficiently infer latent cell-cycle trajectories from scRNA-seq gene expression data. It also can be applied to removing confounding cell-cycle effects, improving classification of cell subpopulations, and enhancing discovery of functional gene subsets. These features make Cyclum useful to constructing the Human Cell Atlas, the Human Tumor Atlas, and other cell ontology.

## Methods

### The Cyclum

The objective of Cyclum is to infer pseudo-time/embedding *x*_*n*_ for cell *n* from its transcriptome profile *y*_*n*_, a column vector containing the expression levels of G genes. Linear methods, such as PCA, find a linear transformation *x*_*n*_ = ℱ (*y*_*n*_) = *Wy*_*n*_ and an inverse linear transformation 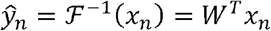, such that the total error 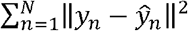 is minimized [34]. Cyclum follows similar formulations, except that the transformation functions ℱ^−1^ (·) and ℱ (·) are nonlinear periodic functions, which makes Cyclum sensitive to circular trajectories (Fig. 1).

We use AutoEncoder [34], a machine learning approach to realize this nonlinear transformation (see Additional file 1, Supplement Text). Specifically, we adopt an asymmetric AutoEncoder (Additional file 1, Figure S1a). In the encoder, we use a standard multi-layer perceptron with tanh(·) activation functions. In the decoder, we use cos(·) and sin(·) as the activation functions in the first layer, followed by a second layer performing linear transformations. These transformations can be represented mathematically as

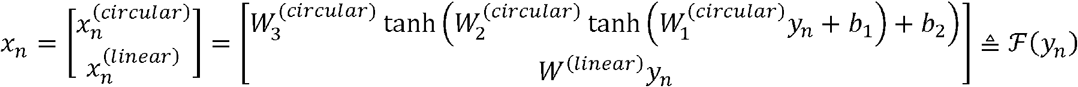

and

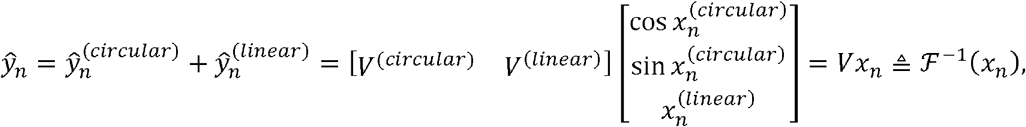

where *W*’s and *b*’s are the weight matrices and translation vectors of the encoder, and *V* is the weight matrix of the decoder. The encoder part is useful when there are a large number of cells. Data from these cells can be divided into mini batches and subsequently loaded into the memory to train the parameters.

We use the least square error as the optimization target with L2 regularization, formally

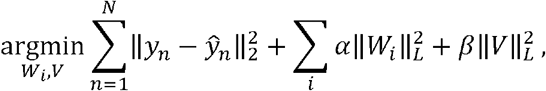

where *W*_*i*_ refers to all the *W*’s presented above. The network is implemented using TensorFlow [33], and the parameters are estimated by gradient descending, using Adam Optimizer. We take the modulus of 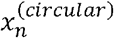 to confine its range to [0,2 π] after the optimization.

### Removing cell-cycle factor

We assume that cell-cycle has an additive effect on the (log-transformed) expression. The 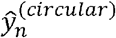 is the estimated cell-cycle effect in *y*_*n*_ and can be removed through subtraction. We then perform t-SNE on the resulting expression levels. For comparison, we use principal components to removing cell-cycle factor by back-transferring the designated principal component to the expression space, subtractinh it from the expression levels. Seurat uses a linear model to find the relationship between gene expression levels and the S and G2M scores it assigns to each cell. The residuals are the expression levels with the cell-cycle factor removed. ccRemover uses a linear GPLVM as the backend to iteratively remove all factors correlated with given cell-cycle genes.

### Predicting marker genes

Using a standard trigonometric identity, the cell-cycle factor of a gene *g* in cell *n* can be reformulated as

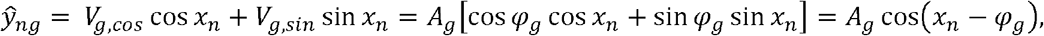

where *φ*_*g*_ is the peak timing of a gene *g* and *A*_*g*_ is the magnitude of the peak, determined by [*V*_*g,cos*_,*V*_*g,sin*_], the *g*’th row of matrix *V*^*(circular)*^. This is an alternative view of the decoder matrix *V*. It means that the decoder assigns pseudo-time to each cell and gives each gene a peak timing and a peak magnitude (Additional file 1,: Figure S1b, c, d). The weight *A*_*g*_ indicates the prominence of the circular pattern in gene *g*. We predict that the important genes are those with higher *A*_*g*_.

### Preprocessing

We used log2 transformed Transcripts Per Million (TPM) in our experiment for scRNA-seq data (the mESC and the melanoma data). For qPCR data (the cell lines) we used, as-is, the reported normalized log counts. One should expect only a slight difference across count normalization methods, as Cyclum examines overall circular patterns, instead of specific values. We also did not filter out any genes or cells, as Cyclum is robust against noise. Standardization was performed on each gene, adjusting the mean expression to 0 and standard deviation to 1 for each gene, so that Cyclum equally considered all the genes.

### Simulating virtual tumor data

To simulate the second clone in the virtual tumor data, we randomly selected a set of cells from the first clone (i.e., the mESC data). We then randomly selected a set of genes from a list of 892 known cell-cycle genes (Additional File 2), and another set from other genes—specifically those that may be affected by, but are not closely related to the cell cycle. We then doubled the expression levels of the selected genes in the selected cells. We also varied the number of cells and genes to simulate data collected from a variety of conditions.

### Metrics

#### Accuracy of timings and separability of subclones

For Cyclone, which outputs categorical cell-cycle phases for each cell, the accuracy is defined as the ratio of cells that are correctly classified. For PCA and reCAT, which output numerical embeddings (pseudo-times), the score is the precision of a best three-component GMM classifier on the embedding [35]. We further assessed the accuracy of the inferred cell-cycle pseudo-time using GSEA against the GO:0007049 GO_CELL_CYCLE signature [17,18], treating pseudo-time as a continuous phenotype and reporting the normalized enrichment score (NES) as the accuracy.

The separability of subclones is defined as the precision of a best two-component GMM classifier on the tSNE of the data. Labels known from independent experiments (i.e., flow-sorting or simulation) are used to evaluate the classifiers.

## Declarations

### Ethics approval and consent to participate

Not applicable in this study.

### Consent for publication

Not applicable in this study.

### Availability of data and material

The source code of Cyclum is freely available for academic use at https://github.com/KChen-lab/cyclum. The datasets generated or analyzed during this study are also available in the repository found at this online location.

## Acknowledgement

This work was supported by a Chan-Zuckerberg Initiative award to Ken Chen.

## Authors’ contributions

SL, FW, JH, and KC developed the Cyclum algorithm. SL implemented the code. SL, FW, JH and KC designed the experiments and helped analyze the results. All authors helped to write the manuscript. All authors have read and approved this paper.

