## Additional File 1 for "Latent periodic process inference from single-cell RNA-seq data"

### Supplementary Text

#### Technical backgrounds

#### Linear ordinary differential equations

Considering the feedbacks and interplays of genes, their expressions may be modeled using linear differential equations [1]. For a single factor (pseudo-time, e.g., cell cycle, senescence), the equation can be written as

$$\frac{d\vec{f}\left( t \right)}{dt}=A\vec{f}\left( t \right),$$

where $\vec{f}(t)$ is a vector that varies over time. Each entry is the expressions of a genes, formally

$$\vec{f}\left( t \right)=\left[ \begin{aligned} \begin{aligned} f_{1}\left( t \right) \\ f_{2}\left( t \right) \\ \vdots\end{aligned} \\ f_{g}(t) \end{aligned} \right].$$

This means the increasing/decreasing rate of gene expressions, are decided by the combination of current gene expressions. The solution is generally a combination of sinusoidal and exponential functions. If we assume the only factor affecting the gene expressions is cell cycle, we expect the solution to be a sinusoidal function. For example, formula

$$\left[ \begin{aligned} f_{1}^{'}\left( t \right) \\ f_{2}^{'}(t) \end{aligned} \right]=\left[ \begin{matrix} 0 & 1 \\ -1 & 0 \end{matrix} \right]\left[ \begin{aligned} f_{1}\left( t \right) \\ f_{2}(t) \end{aligned} \right]$$

means $f_{1}$ promotes transcription of $f_{2}$, while $f_{2}$ inhibits transcription of $f_{1}$. Notably, $\left[ \begin{aligned} f_{1}\left( t \right) \\ f_{2}(t) \end{aligned} \right]=\left[ \begin{aligned} \sin t \\ \cos t \end{aligned} \right]$ is one of the solutions, which means this kind of regulation will result in a circular process. This explains why sinusoidal function is a good way to model the genes.

##### Partial differential equation

There is a significant difference between real-time and pseudo-time. For each cell, the real-time is always unidimensional. However, for trajectory inference, it is important to remember that the cells on the trajectory are not really a single cell changing over time. Having more time axes (e.g., cell cycle and differentiation) allows for better modeling. In general cases, several processes may jointly affect the gene expression, introducing multiple “timers” dictating the cell fate, resulting in a partial differential equation. Nevertheless, we may mimic the separation of variables method and assume that the (log-transformed) gene expressions can be separated into addable terms and still use ordinary differential equation components to find the solution for each term. The combination of linear and non-linear kernels is a realization of this idea (i.e. the summation of first and second order differential equations).

For example, we may have a partial differential equation with two time factors, $s$ and $t$:

$$\frac{\partial\vec{f}\left( s,t \right)}{\partial s}+\frac{\partial\vec{f}\left( s,t \right)}{\partial t}=A\vec{f}\left( s,t \right).$$

We further assume that $\vec{f}\left( s,t \right)=\vec{S}\left( s \right)+\vec{T}(t)$, where $\vec{S}\left( s \right)$ and $\vec{T}(t)$ only depend on $s$ and $t$, respectively. This uses the “separation of variables.” Then, we have

$$\frac{\partial\vec{S}\left( s \right)}{\partial s}+\frac{\partial\vec{T}\left( t \right)}{\partial t}=A\left( \vec{S}\left( s \right)+\vec{T}\left( t \right) \right),$$

which can be solved for $s$ and $t$ individually. In other words, it can be written into two ordinary differential equations. This is roughly the background of having more than one dimension of pseudo-time in the Cyclum embedding.

##### Deconvolution

Ordinary differential equations can be depicted as a linear system. Cyclum infers pseudo-time over which the expression can be explained by Fourier expansion truncated to the first term. From the perspective of frequency domain, Cyclum eliminates that term to remove the cell-cycle factor. According to time-frequency duality, from the perspective of time domain, this is a deconvolution. The cell-cycle factors are removed by the deconvolution.

### Supplementary Figures

a

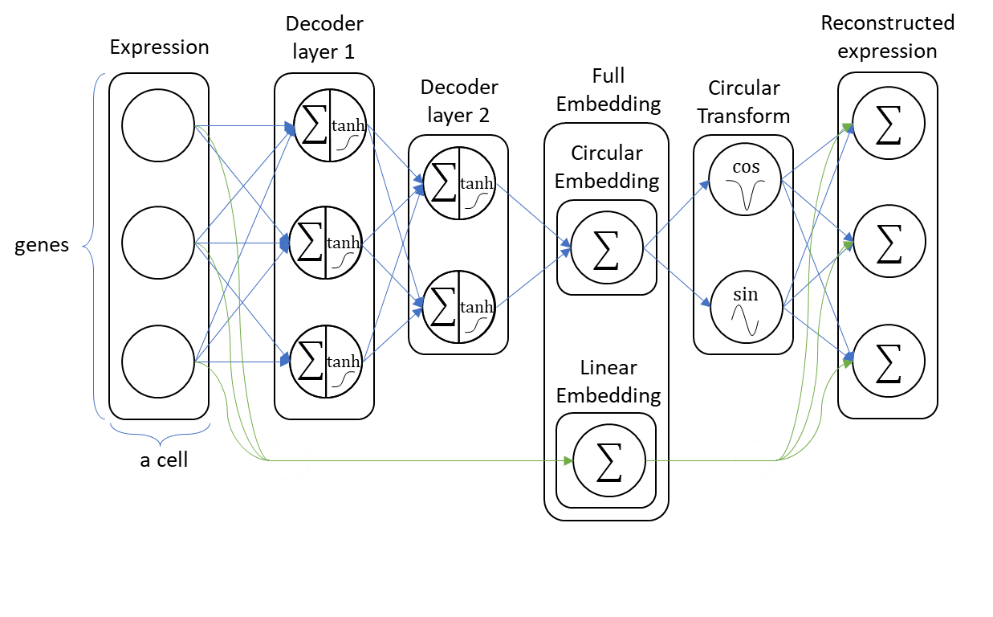

b

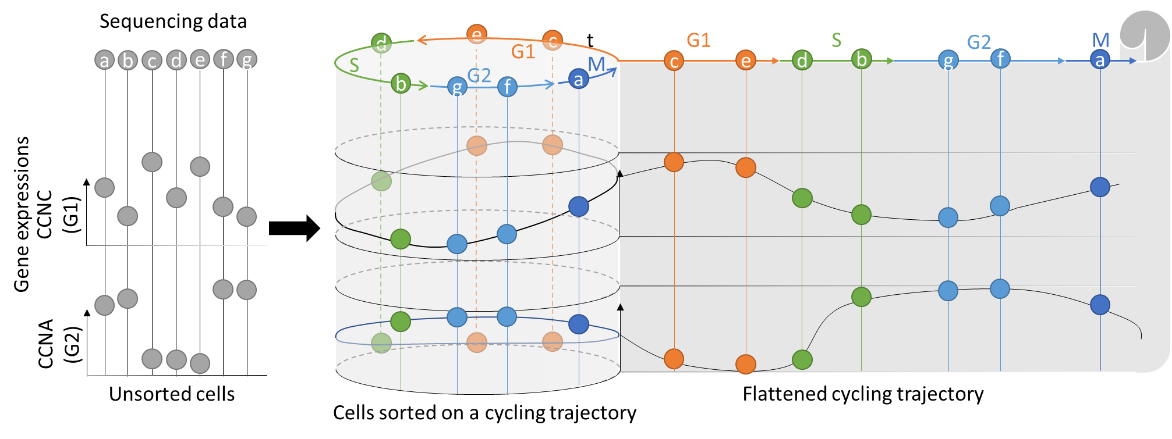

c	d

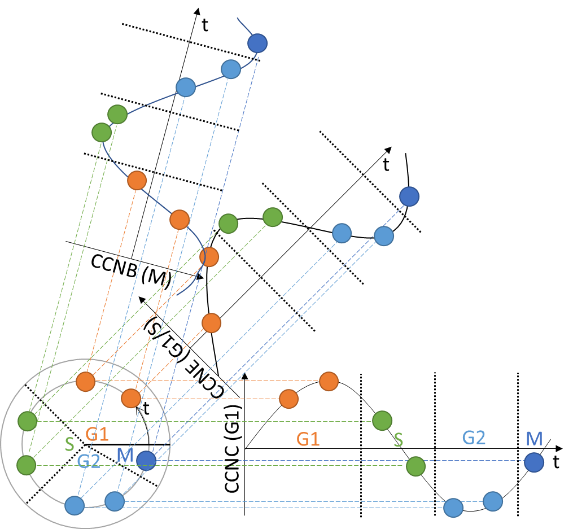

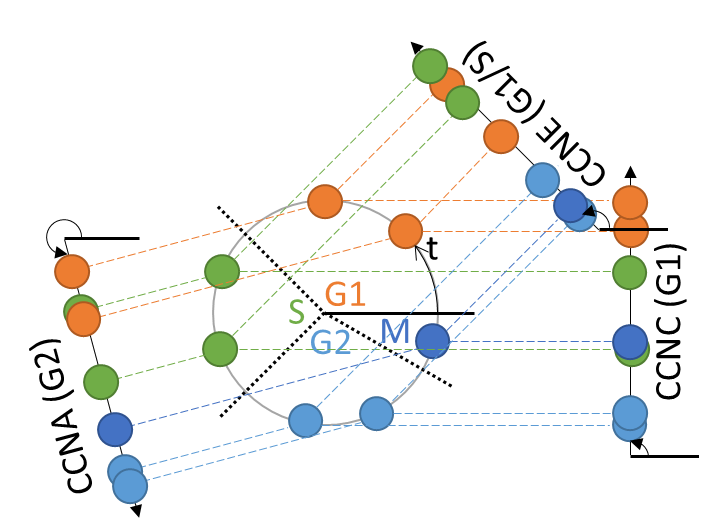

**Figure S1:** Mechanism of Cyclum. **a)** The diagram of the auto-encoder. **b)** The unravel of the periodic pseudo-time. **c)** The optimal sinusoidal representation of Cyclum. **d)** The optimal sinusoidal representation of Cyclum, without showing timings for genes.

In the sequencing data, cells are unsynchronized. Cyclum infers a real-value timing “coordinate” for each cell on a circle. The expressions of genes are assumed to be periodic and follow a roughly sinusoidal pattern through the time. The gene expression can be viewed as the projection of the cycle onto an axis. The angle of the axis is the phase of the gene (i.e. peak timing in radian measure).

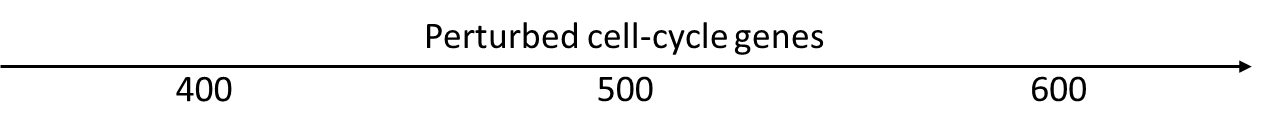

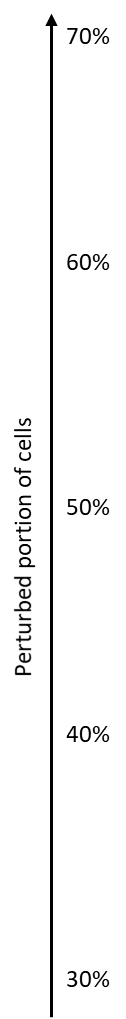

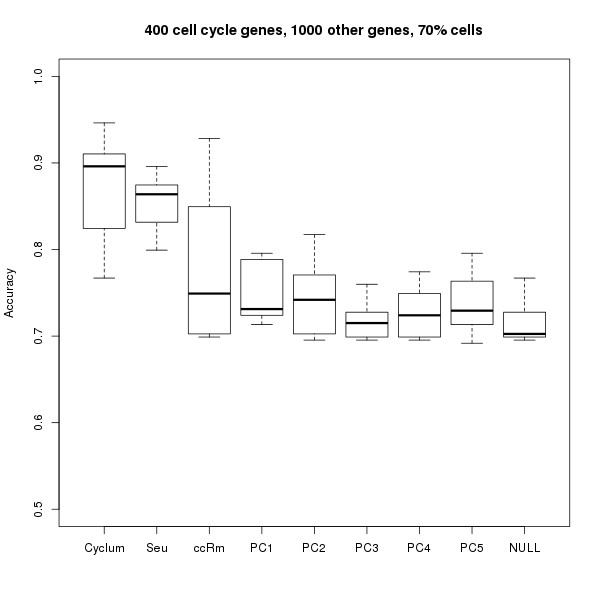

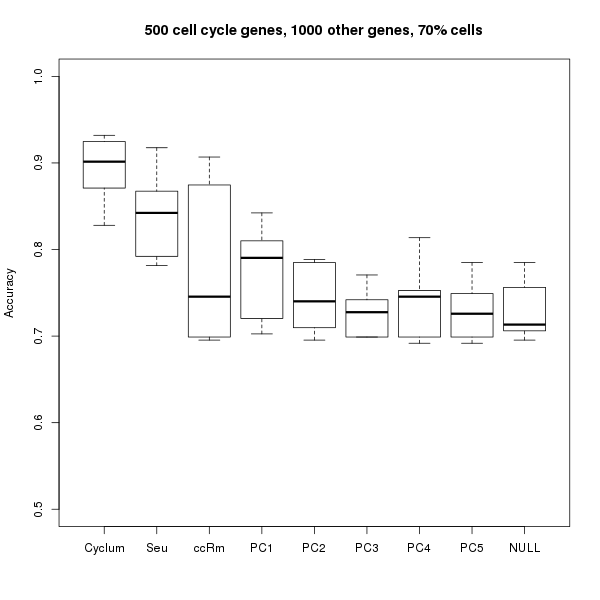

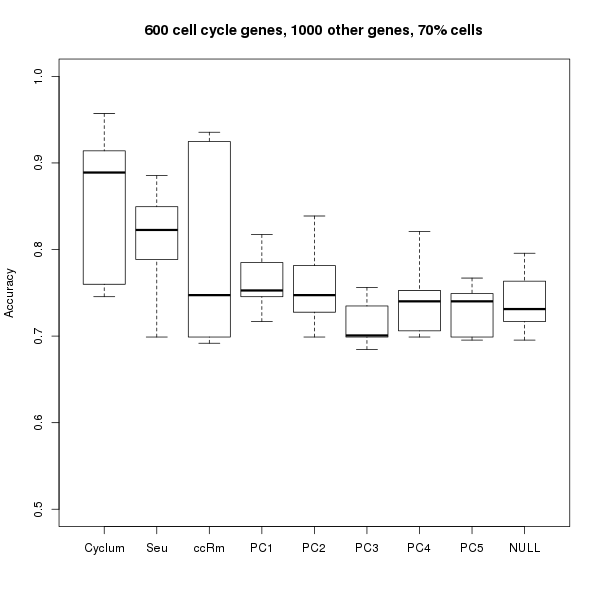

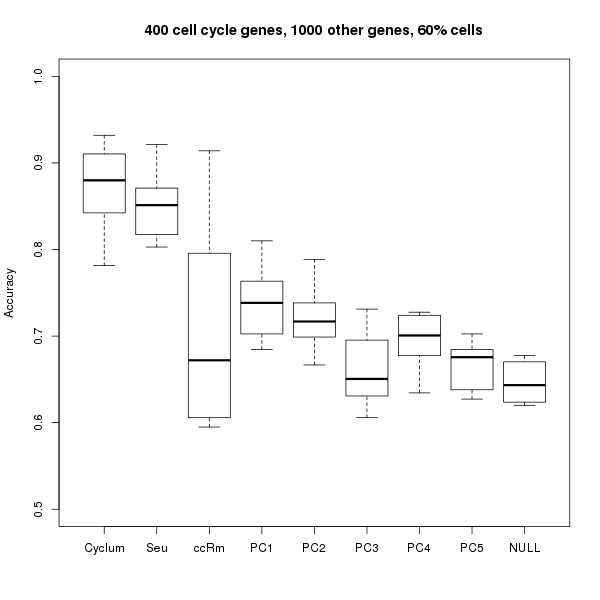

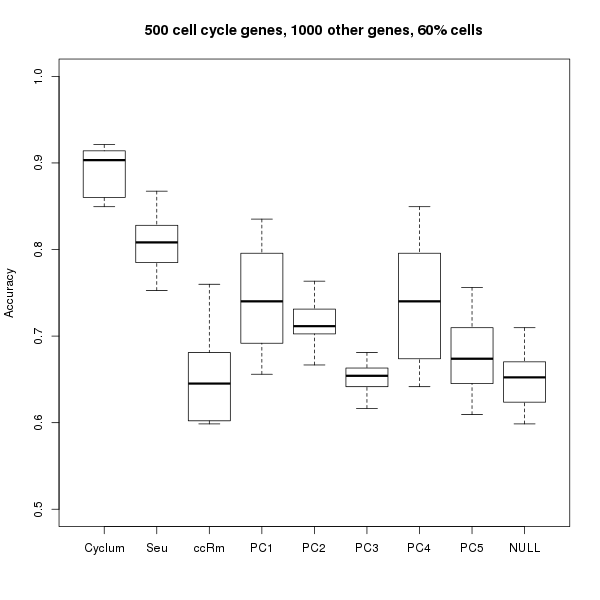

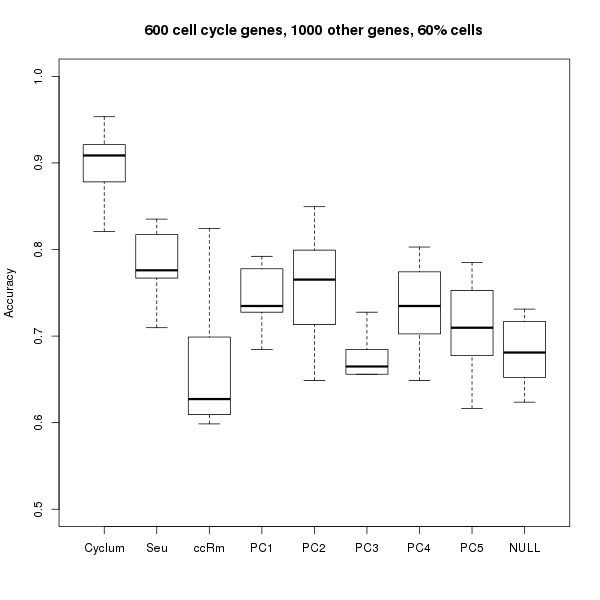

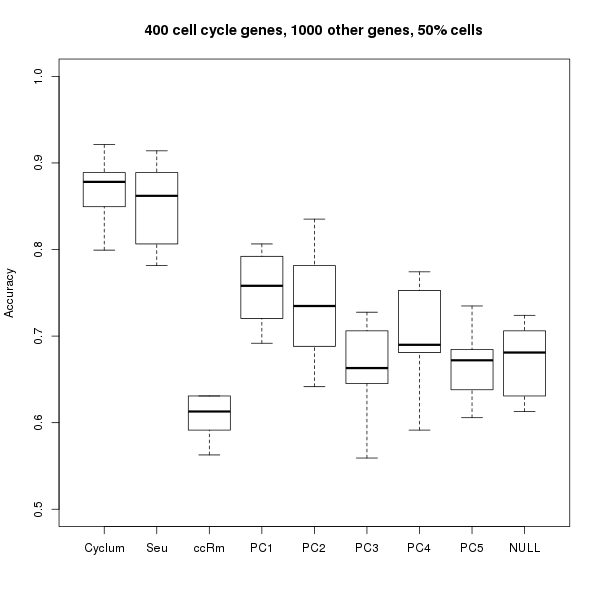

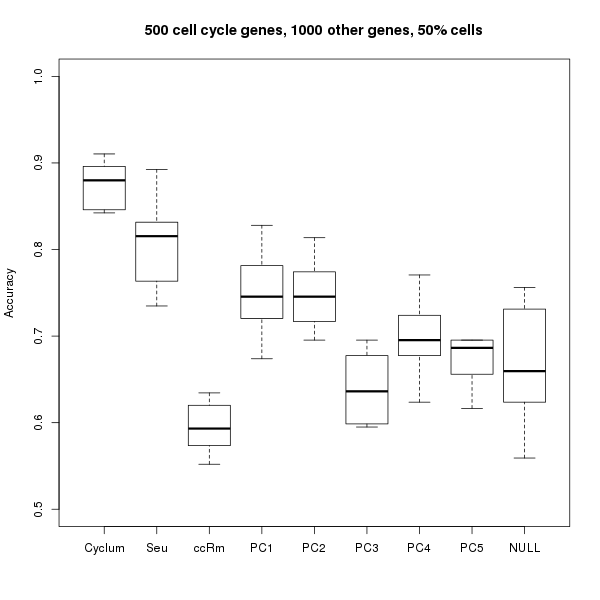

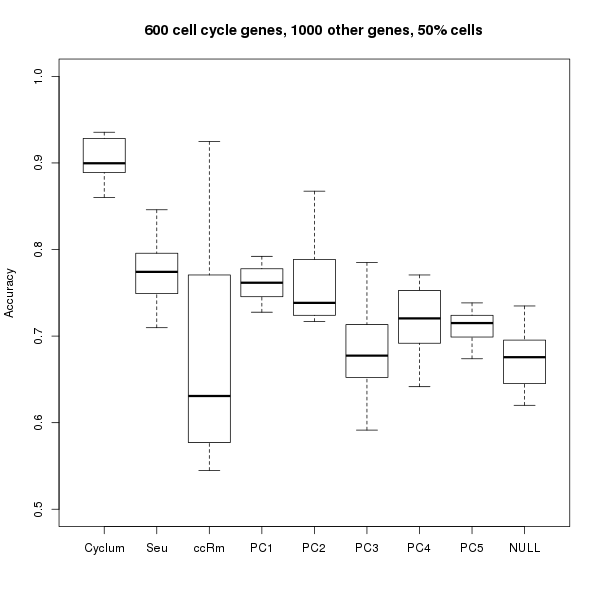

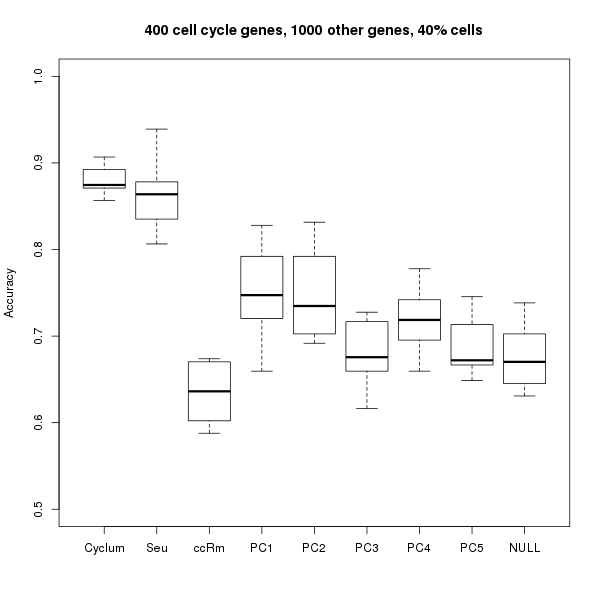

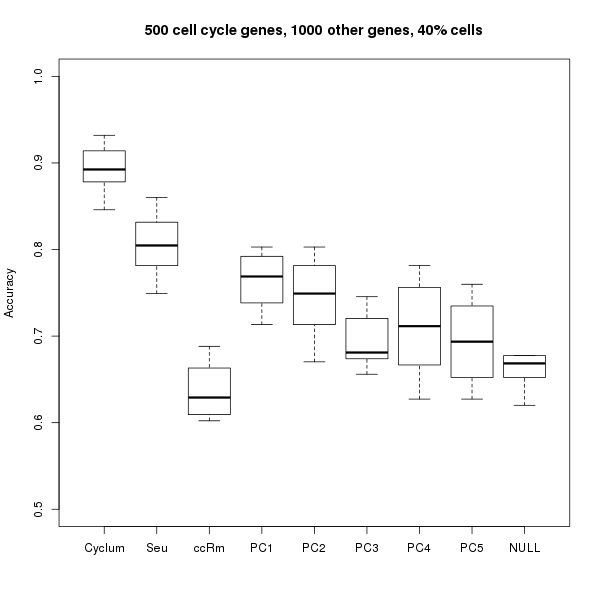

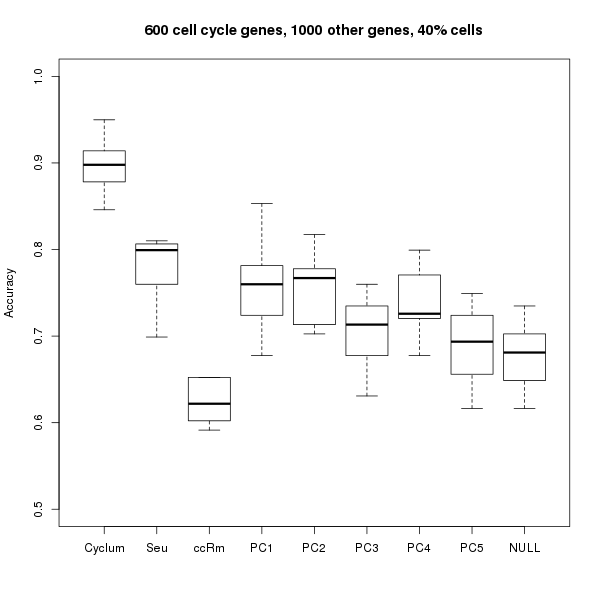

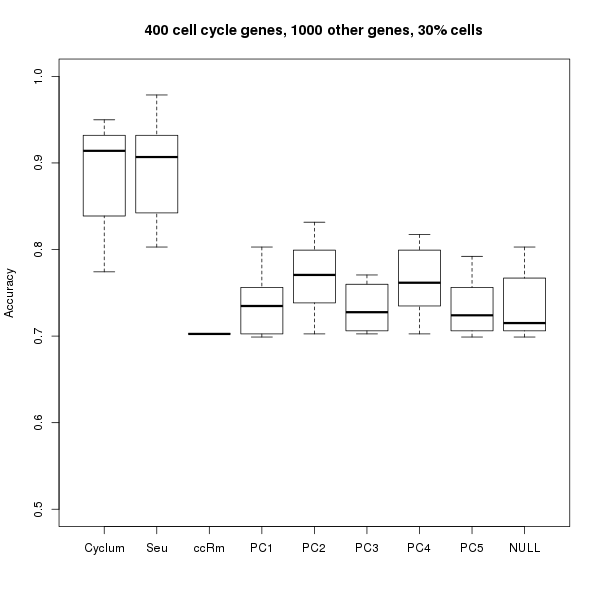

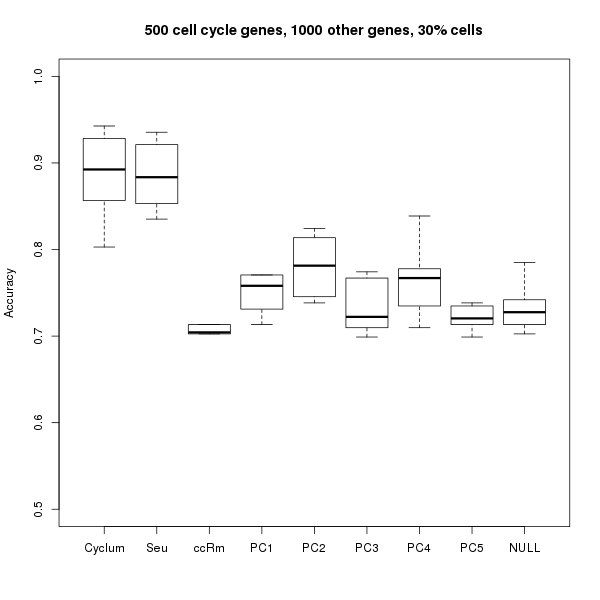

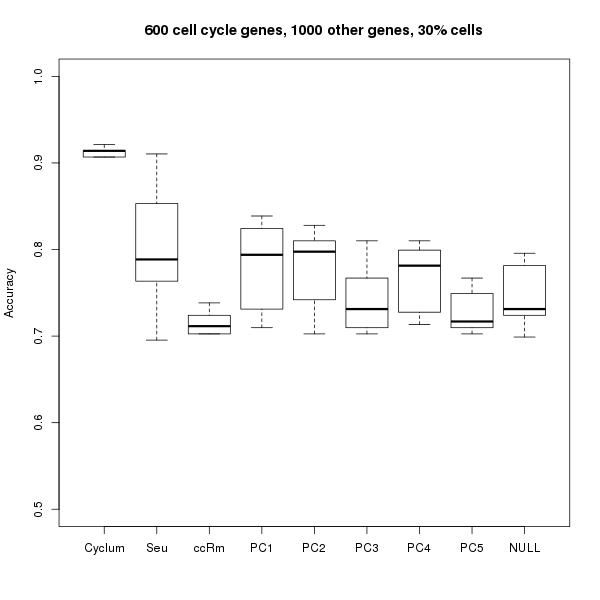

**Figure S2:** Separability of subclones in more experimental settings varying number of perturbed cell cycle genes and portion of perturbed cells. Each box plot show the range (box) and the mean (horizontal bar) separablility (Y-axis) of 10 randomly generated virtual tumor datasets using data corrected by Cyclum, Seurat (Seu), ccRemover (ccRm), and Principal Component (PC) 1-5, and the uncorrected data (NULL). From left to right, the columns are for perturbing 400, 500, and 600 cell-cycle genes, repectively. The number of other genes perturbed is fixed to 1000. From bottom to top, the rows are for perturbing 30%, 40%, 50%, 60%, and 70% of the cells to form a new subclone.

We compare the three experiments perturbing different number of genes directly related to cell-cycle and portion of cells, while the number of other genes perturbed is fixed to 1000. When less cell-cycle genes are perturbed (first column), the performance of Cyclum is similar to Seurat. However, Cyclum performs significantly better when more cell-cycle genes are perturbed (second and third columns). Model-based construction leads to this improvement. For ccRemover, the performance is not robust. For PCA, the best PC to remove shifts, showing that this is not a easy problem to solve using this method.

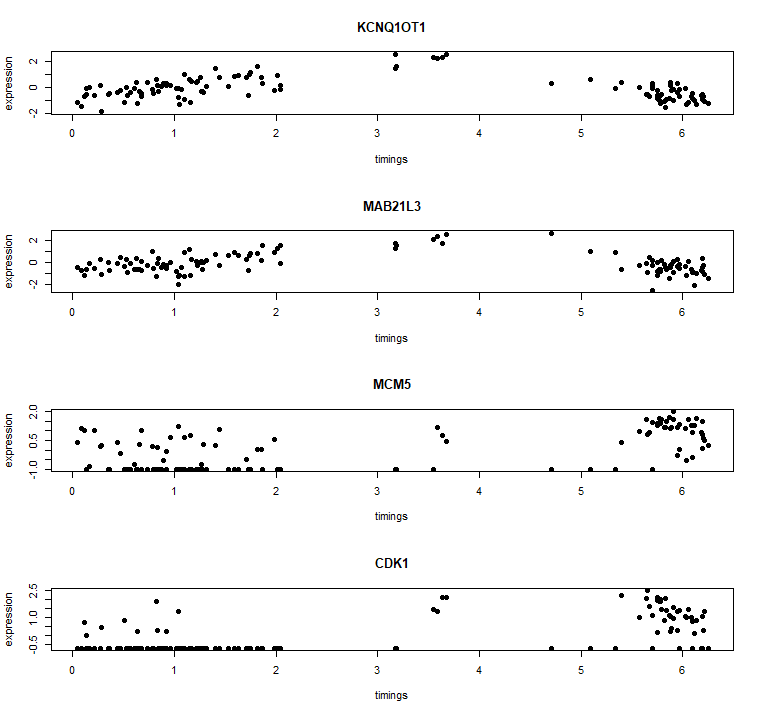

**Figure S3:** Top genes found by Cyclum compared with known cell cycle genes. These plots show the correlation of the expressions (Y-axis) of the top genes found with inferred pseudo-time (X-axis).

Two of the top genes reach their maxima differently from known cell cycle genes MCM5 and CDK1. This is a unique advantage of Cyclum. The non-coding RNA KCNQ1OT1 shows up frequently in recent studies [2–5], linking it to (malignant) cell proliferation, although its role is still debatable. As another example, recent studies also show FBLIM1 as a marker of malignancy [6,7]. Other top genes found by Cyclum are shown in Table S2.

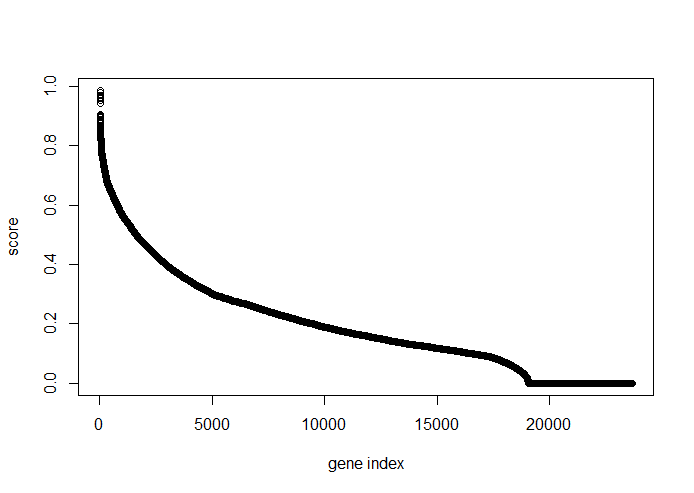

**Figure S4:** Cyclum assigned cycling score for the genes. A gene with the rank of x, shown on the X-axis, has the score y, shown on the Y-axis.

Cyclum assigns a cycling score for the genes. The distribution of the score is not flat. There are some genes with dramatically higher scores, while the majority of genes have significantly lower scores.

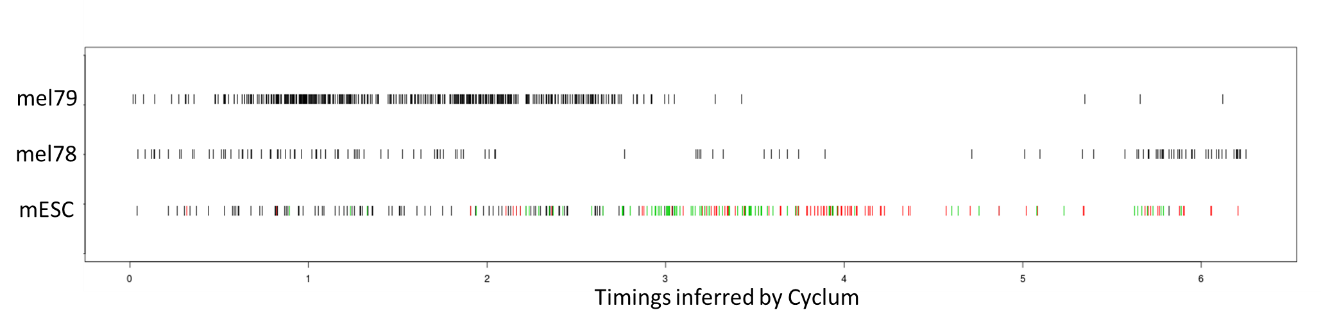

**Figure S5:** Comparison of pseudo-times. This chart shows the comparison of pseudo-times (X-axis) inferred on mESC, mel78, and mel79 (Y-axis). Stages of mESC cells (active cycling) are encoded in color (Black: G0/G1; green: G2/M; red: S). Stages of mel78 (partially cycling) cells and mel79 (almost non-cycling) cells are unknown.

For actively cycling mESC cells, the distribution of cells on the pseudo-time axis is relatively uniform. Mel78 contains about 40% cycling cells, most cells cluster in time (0, 2), which may be G0/G1. G2/M may be in (3, 4) and S may be in (5, $2\pi$). For Mel79, which reportedly is not under active cycling, almost all cells are clustered between (0, 3) and the interval (3, $2\pi$) is almost vacated. Although the pseudo-time cannot be precisely compared across datasets, it provides a quantitative way to measure the portion of cells in each cycling stage.

a	b	c	d

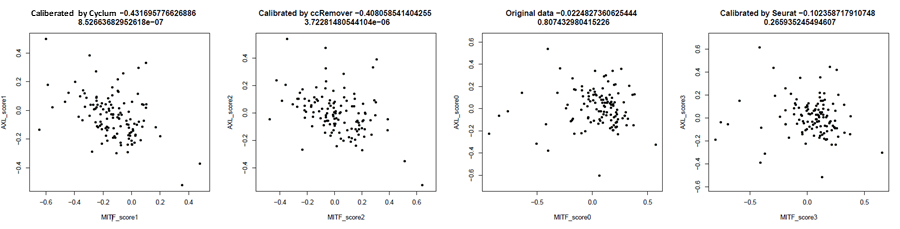

e	f

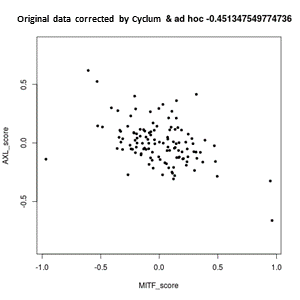

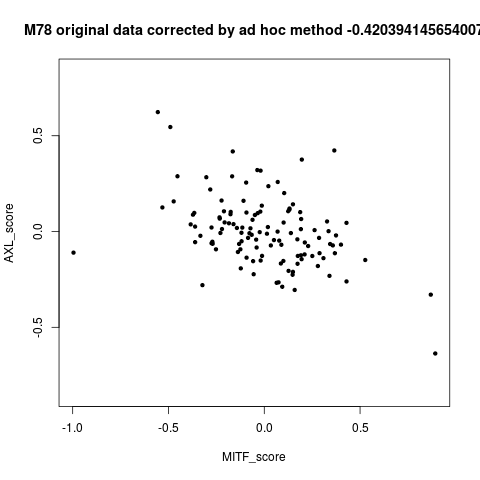

**Figure S6:** AXL and MITF program. For a-d, the correlation of MITF score (X-axis) and AXL score (Y-axis) on expressions corrected by **a)** Cyclum, **b)** ccRemover, and **d)** Seurat, compared with **c)** the uncorrected original expressions. For e and f, the expressions corrected by **e)** both Cyclum and the ad hoc method. **f)** only the ad hoc method suggested in the original report.

Cyclum yields the best negative correlation, as compared to ccRemover, Seurat, and uncorrected data. Combining Cyclum with the ad hoc approach applied in the original paper also improves the negative correlation.

| **Table S1: The melanoma datasets** | | | | |
| --- | --- | --- | --- | --- |
| Patient | Number of cells | Number of malignant cells | Percentage of cycling cells in the report | Percentage of cycling cells inferred using Cyclum |
| 79 | 896 | 468 | 1% | 3% |
| 81 | 205 | 133 | (likely low cycling) | 11% |
| 80 | 480 | 125 | (likely high cycling) | 60% |
| 78 | 131 | 120 | 29% | 42% |
| 88 | 351 | 117 | (likely high cycling) | 7% |

For Mel80, 81, and 88, percentages are not reported, but the Figure S4C in the original report suggests whether or not they are high cycling.

| **Table S2: Enrichment AUC on Melanoma** | | | |
| --- | --- | --- | --- |
| Patient | Cyclum | PC 1 | PC 2 |
| 79 | **0.641** | 0.594 | 0.529 |
| 81 | 0.602 | **0.616** | 0.565 |
| 80 | **0.649** | 0.639 | 0.569 |
| 78 | **0.649** | 0.626 | 0.623 |
| 88 | **0.585** | 0.543 | 0.527 |

We count the numbers of known cell-cycle genes in the top $G$ proposed genes, where $G$ is an integer. We vary $G$ from one to the number of genes. The sum of resulting number of cell-cycle genes over the $G$’s is the enrichment AUC.

| **Table S3: The proposed top genes** | | | |
| --- | --- | --- | --- |
| Gene | Weight | Gene | Weight |
| KCNQ1OT1 | 0.986 | COPS8 | 0.813 |
| MAB21L3 | 0.979 | LSM10 | 0.812 |
| LOC100131257 | 0.970 | ARF4 | 0.810 |
| UGDH-AS1 | 0.960 | BLVRB | 0.809 |
| FBLIM1 | 0.960 | PGAM1 | 0.807 |
| ORC4 | 0.954 | PSMA2 | 0.806 |
| ABCC9 | 0.941 | PKMYT1 | 0.804 |
| TLCD2 | 0.904 | NDUFA8 | 0.804 |
| SF3B14 | 0.902 | TPTE2P1 | 0.800 |
| REXO1L1 | 0.899 | ASTN2 | 0.800 |
| C21orf62 | 0.893 | COX7A2L | 0.800 |
| HYDIN2 | 0.889 | S100B | 0.798 |
| ATP5G3 | 0.885 | NDUFB10 | 0.795 |
| LOC646214 | 0.879 | PSMD8 | 0.793 |
| FTH1 | 0.878 | LINC00346 | 0.790 |
| ANKRD20A9P | 0.878 | ADIPOR1 | 0.786 |
| RPS3 | 0.878 | ATOX1 | 0.785 |
| PRKAR1A | 0.876 | SLC25A3 | 0.785 |
| SLC25A6 | 0.866 | KLRD1 | 0.783 |
| SHISA9 | 0.866 | MTRNR2L10 | 0.783 |
| PSMB4 | 0.861 | SRP72 | 0.782 |
| FABP5 | 0.859 | TXN | 0.781 |
| NME1 | 0.857 | PYCARD | 0.780 |
| LOC643406 | 0.855 | NDUFS7 | 0.779 |
| LAMTOR1 | 0.850 | NDUFB3 | 0.778 |
| GSTO1 | 0.849 | EIF5A | 0.778 |
| MTRNR2L4 | 0.844 | TRMT112 | 0.778 |
| TSTD1 | 0.843 | SOD1 | 0.777 |
| S100A11 | 0.842 | MIF | 0.777 |
| ATCAY | 0.842 | MS4A10 | 0.775 |
| NLRP12 | 0.836 | KPNA2 | 0.775 |
| ATP5G1 | 0.836 | HADHB | 0.774 |
| TSIX | 0.835 | ROMO1 | 0.774 |
| RPL3 | 0.835 | ALG8 | 0.773 |
| ARHGEF26-AS1 | 0.832 | MLANA | 0.773 |
| CHMP2A | 0.831 | ZBTB8A | 0.772 |
| COA4 | 0.830 | SDCBP | 0.772 |
| COX6C | 0.829 | TUBB | 0.772 |
| RPL29 | 0.826 | LGALS3 | 0.770 |
| GSTP1 | 0.826 | CCBE1 | 0.770 |
| IKZF3 | 0.824 | HNRNPM | 0.769 |
| SPN | 0.824 | MTRNR2L6 | 0.768 |
| PDDC1 | 0.823 | ANXA5 | 0.768 |
| DBI | 0.820 | PARK7 | 0.768 |
| ODF2L | 0.819 | UHRF1 | 0.768 |
| ATP5A1 | 0.819 | CD84 | 0.767 |
| COX7A2 | 0.819 | SNRPE | 0.767 |
| C1QBP | 0.818 | PAICS | 0.767 |
| ARGFX | 0.815 | COX6B1 | 0.767 |
| OST4 | 0.815 | RPL8 | 0.767 |
